# Optimised cell systems for the investigation of hepatitis C virus E1E2 glycoproteins

**DOI:** 10.1101/2020.06.18.159442

**Authors:** Mphatso D. Kalemera, Joan Cappella-Pujol, Ana Chumbe, Alexander Underwood, Rowena A. Bull, Janke Schinkel, Kwinten Sliepen, Joe Grove

## Abstract

Great strides have been made in understanding and treating Hepatitis C virus (HCV) thanks in part to the development of the full-length cell-culture system, the pseudoparticle system and soluble envelope glycoproteins. The HCV pseudoparticle (HCVpp) system is a platform used extensively in studies of cell entry, screening of novel entry inhibitors, assessing the phenotypes of clinically observed E1 and E2 glycoproteins and, most pertinently, in characterising neutralizing antibody breadth induced upon vaccination and natural infection in patients. Nonetheless, some patient-derived clones fail to produce infectious particles or produce particles that exhibit infectivity too low for meaningful phenotyping. The mechanisms governing whether any particular clone produces infectious pseudoparticles are poorly understood. Here we show that endogenous expression of CD81, an HCV receptor and a cognate binding partner of E2, in producer HEK 293T cells is detrimental to the infectivity of recovered HCVpp for most strains. Many HCVpp clones exhibited increased infectivity or had their infectivity rescued when they were produced in HEK 293T cells CRISPR/Cas9 engineered to ablate CD81 expression (293T^CD81KO^). Clones made in 293T^CD81KO^ cells were antigenically very similar to their matched counterparts made parental cells and appear to honour the accepted HCV entry pathway. Deletion of CD81 did not appreciably increase the recovered titres of soluble E2 (sE2). However, we did, unexpectedly, find that monomeric sE2 made in 293T and 293F exhibit important differences. We found that 293F-produced sE2 harbours mostly complex type glycans whilst 293T-produced sE2 displays a heterogeneous mixture of both complex type glycans and high-mannose or hybrid type glycans. Moreover, sE2 produced in HEK 293T cells is antigenically superior; exhibiting increased binding to conformational antibodies and the large extracellular loop of CD81. In summary, this work describes an optimal cell line for the production of HCVpp and reveals that sE2 made in 293T and 293F cells are not antigenic equals. Our findings have implications for functional studies of E1E2 and the production of candidate immunogens.

## Introduction

Hepatitis C virus is a significant human pathogen infecting more than 70 million people worldwide, of whom the majority are chronically infected. Current WHO estimates suggest around half a million patients succumb to the disease annually, mostly due to complications arising from cirrhosis or hepatocellular carcinoma. Transmission still continues unabated with incidence rates rising in North America as the majority of infected individuals are unaware of their status (1). The recent development of curative direct-acting antivirals (DAAs) has revolutionized HCV therapy as well as raised the possibility of eliminating HCV. Nonetheless, the high cost of treatment, the risk of reinfection following successful treatment and poor awareness of HCV status in high-risk groups necessitate the development of a prophylactic vaccine (2).

As viruses are genetically diverse it is imperative that experimental systems have the capacity to include many isolates allowing for comprehensive assessment of the effectiveness of therapeutic and prophylactic interventions (3–5). There are two major systems to assess HCV infection in vitro: HCV pseudoparticles (HCVpp) and cell-culture derived HCV (HCVcc). HCVpp are based on a disabled retroviral construct, which encodes a reporter gene and incorporates the HCV glycoproteins E1 and E2 in their lipid envelope (6,7). HCVcc are full-length replicative viruses generated by the introduction of in vitro transcribed RNA genome in to permissive cells; HCVcc are typically based on the JFH-1 clone or chimeras consisting of the JFH-1 replicase genes NS3-NS5B and Core-NS2 regions of alternative HCV genomes (8–10).

The HCVcc system represents a more physiological model of infection and allows for a broader study of the HCV life cycle (11). However, this system still largely depends on JFH-1 chimeras, which often rely on culture adaptation for optimal infectivity, and has challenging production and handling protocols given its biosafety level III designation in most regions (12). In contrast, producing HCVpp is a relatively simple task and the ease of handling allows for the simultaneous generation of a multitude of clones if one requires (13). The major disadvantage of HCVpp is that their application is only limited to studying viral entry. Regardless, due to its flexibility, the HCVpp system is preferred for characterising neutralizing antibody breadth, an essential component for screening HCV vaccine candidates. To this end, Wasilewski and colleagues recently observed a very strong positive correlation between the relative neutralization resistance of E1E2-matched HCVcc and HCVpp variants indicating that either system can be used to phenotype neutralizing antibodies (14). Furthermore, diverse HCVpp panels were key in identifying the relationship between the development of broadly neutralizing antibodies and spontaneous clearance of HCV infection (15–17).

Another important tool for studying HCV entry and vaccine development is a soluble form of its glycoprotein E2 (sE2), which is devoid of its transmembrane region and is expressed in the absence of E1. Screenings based on sE2 binding to human hepatoma cell lines identified the first two HCV entry receptors as being the tetraspanin CD81 and scavenger receptor class type B 1 (SR-B1) (18,19). sE2 retains proper folding as evidenced by its ability to recapitulate HCV binding to CD81 and SR-B1, block HCVcc infection and bind various antibodies (20), therefore allowing for functional, structural and biophysical characterisation of E2. The partial structures of sE2 revealed a globular shape with IgG-like folds and flexible regions (21,22). Interestingly, computational models and antibody competition studies suggest there may be an interdependence in E2-SR-B1/CD81 interactions, whereby SR-B1 binding enhances E2-CD81 engagement (23,24). The SR-B1 and CD81 binding epitopes on E2 are predicted to be in close proximity thus steric clashes may prevent simultaneous engagement of the two receptors. However, several studies have recently suggested this region may be highly flexible thereby potentially introducing a greater spatial separation between the epitopes (25–27). Investigating these hypotheses will further clarify our understanding of HCV entry as well as aid vaccine development and sE2 will doubtless continue to be a key tool to decipher these mechanisms.

HEK 293-derived cell lines are extensively used for the production of many recombinant proteins and pseudoviruses, including sE2 and HCVpp. HEK 293 cells express CD81 (28,29), however it remains unknown whether the presence of this cognate E2 receptor during production affects the functionality and antigenicity of pseudovirus and/or sE2. Here, we describe the generation of a 293T cell line knocked out for CD81 expression (293T^CD81KO^). Producing HCVpp in this cell line significantly improved or rescued the infectivity of the majority of clinical isolates screened (30) without altering particle antigenicity. Finally, we show that while sE2 molecules produced in 293T^CD81KO^ and 293T cells are very similar they are antigenically superior to that produced in 293F cells. We propose this is likely due to observed differences in glycosylation status between the two.

## Results

### HCVpp supernatant produced in CD81 knock-down 293T cells exhibits enhanced infectivity

Our study was initiated following an observation we made during routine experiments: conditioned media taken from 293T cells contains a high level of CD81. Figure 1a displays extracellular CD81 in filtered conditioned cell-culture supernatant from two cell lines extensively used in HCV research (the Huh-7 and 293T cell lines). We detected CD81 in both 293T and Huh-7 conditioned supernatant, although the former contained considerably more (Fig. 1a). The extracellular source of CD81 is likely to be exosomes: small extracellular vesicles formed by the inward budding of the late endosomal membrane. They measure 30-120 nm in size and are composed of proteins, lipids, nucleic acids and other metabolites. Notably, their surfaces are highly enriched in tetraspanins such as CD81, which consequently serves as an exosome biomarker.

**Fig. 1.**
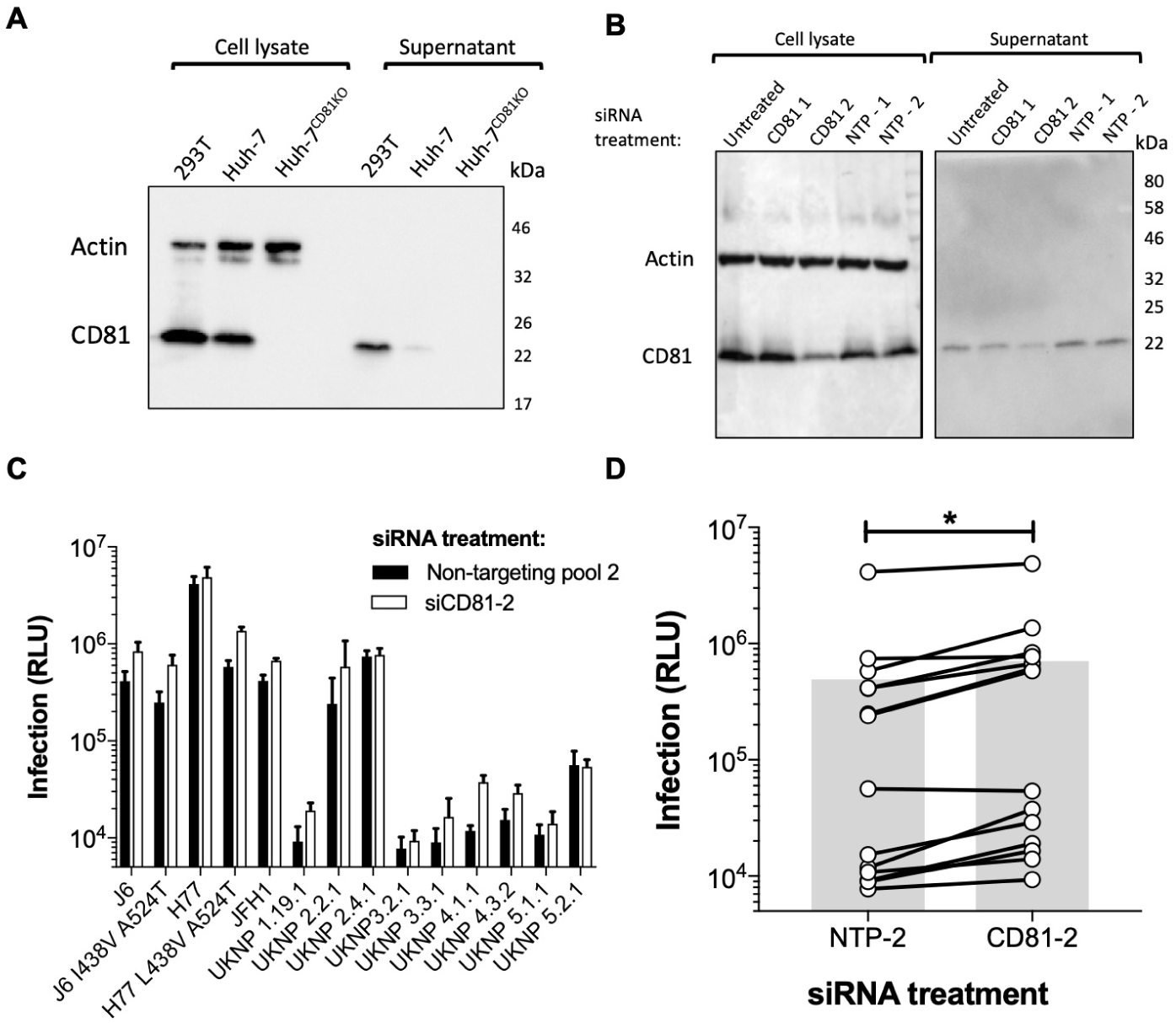
Knockdown of CD81 in producer 293T cells increases HCVpp infectivity. Cell lysates and unconcentrated conditioned supernatant of **(A)** untreated cells and **(B)** CD81 targeting siRNA transfected 293T cells were analysed by SDS-PAGE and Western blot with a specific anti-CD81 antibody. Molecular mass markers are indicated on the right (kDa). **(C)** Huh-7 cells were challenged with an HCVpp panel consisting of 14 E1E2 clones produced in 293T cells exhibiting appreciable CD81 knockdown (black bars) or 293T cells treated with a non-targeting siRNA pool (NTP-2) (white bars). Data are represented as raw luciferase units (RLU) and are from a single experiment performed in triplicate. Error bars indicate the standard deviations between three replicate wells. **(D)** A compiled summary of the data shown in c, connected points indicate a single E1E2 clone and the grey bars represent the mean RLU of all clones. Paired t-test, (*p < 0.05).

We theorised that exosomes and HCV particles may complex extracellularly through interactions between surface CD81 and E2, and that this may be detrimental for HCVpp infectivity. To test this, we screened the infectivity of a panel of HCVpp produced in 293T cells treated with siRNA targeting CD81 (Fig. 1b). We consistently observed greater infection of Huh-7 cells for HCVpp produced in CD81 knockdown 293T cells (CD81-2) compared to their E1E2 matched equivalents produced in non-targeted pool siRNA (NTP-2) treated 293T cells (Fig. 1c). Overall, CD81 knock-down led to a two-fold increase in infection for viruses in the panel, collectively (Fig. 1d) indicating that the expression of CD81 in 293T cells affects HCVpp production. We were unable to demonstrate cell-free interactions between HCVpp produced in CD81 knockdown 293Ts and exosomes (data not shown); however, it is also possible that CD81 affects HCVpp production intracellularly or during viral egress (see discussion).

### Deletion of CD81 in 293T cells enhances or rescues infectivity of patient-derived HCVpp

Following the above observations (Fig. 1d) we decided to generate stable cell lines ablated of CD81 expression as they could be a useful resource for the production of HCVpp (30,31). To delete CD81, we engineered 293T cells with the CRISPR-Cas9 gene-editing system. As CD81 is a cell surface protein we were able to sort edited cells by flow cytometry to obtain pure CD81 negative populations (coloured boxes in Fig. S1a). Sorted cells were then diluted to obtain single cell clones and expanded (Fig. S1b) until we obtained two stable cell lines: 293T^CD81KO^ guide 1 and guide 3 (g1 and g3) (Fig. 2a).

**Fig. 2.**
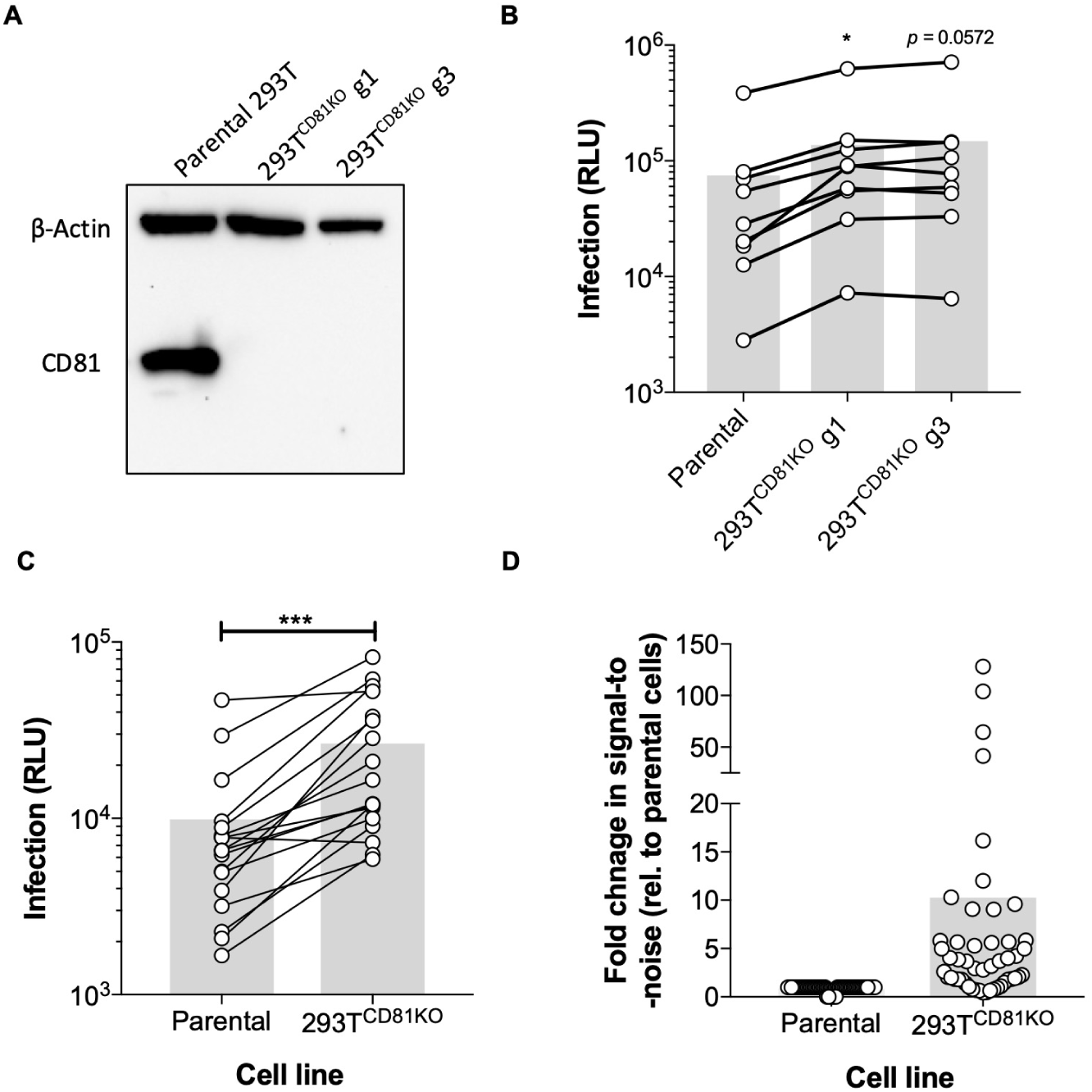
HCVpp made in CD81 knock-out 293T cells exhibit enhanced infectivity. Huh-7 cells were challenged with HCVpp produced in parental or 293T^CD81KO^ cell lines. **(A)** CD81 ablation was confirmed by SDS-PAGE and Western blot analysis. **(B)** A compiled summary of the infection levels for an HCVpp panel consisting of both prototypical and clinical isolates. Paired t-test with virus produced in parental 293Ts (*p < 0.05). **(C)** A compiled summary of the infection levels of a panel of clinical HCVpp isolates. Paired T-test, (***p < 0.001). Data in b and c are from lone experiments performed in triplicate and data are represented as RLU. **(D)** A compiled summary of the fold signal-to-noise ratio (S/N) for all screened clones, fold values are relative to virus produced in parental 293T cells. Connected points indicate a single E1E2 clone and the grey bars represent the mean value.

To comprehensively determine whether CD81 deletion enhanced HCVpp infection, several infection screens comparing the infectivity of HCVpp produced in parental 293Ts to that made in the CD81 knock-out cell lines were performed in three independent labs. Firstly, a panel consisting of both prototypical strains (e.g. J6 and H77) and clinical isolates was made in the aforementioned cell lines. We consistently observed higher infection across the panel for HCVpp harvested from both 293T^CD81KO^ lines (g1 and g3) compared to matched viruses made in parental cells (Fig. 2b & S2a-b). As the infectivities of HCVpp recovered from the two 293T^CD81KO^ cell lines were very similar, we chose the g3 CD81 knock-out lineage for subsequent experiments, it is henceforth referred to as 293T^CD81KO^. The next screen was of a panel of HCVpp carrying patient-derived E1E2 observed in patient cohorts from hospitals in Amsterdam (32 and unpublished) or Nottingham (33). Here, the improvement in infection for HCVpp produced in 293T^CD81KO^ cells compared to their equivalents made in parental cells was markedly more apparent (Fig. 2c & S2c-d). Indeed, we observed a five-fold increase in infectivity for four of the tested clones and this effect reached ten-fold for the UKNP 5.2.1 clone (Fig. S2c).

Lastly, we made a panel of HCVpp expressing E1E2 observed in HCV^+^ individuals from the Australian Hepatitis C Incidence and Transmission Studies in prisons (HITS-p) cohort in the 293T^CD81KO^ cells (34). Following infection readout, we calculated the signal-to-noise ratio (S/N) for each clone in the panel and compared this value to a previously determined S/N when the clone was produced in parental 293Ts (Table S1). We saw an increase in the S/N for the majority of screened clones when made in 293T^CD81KO^ cells. Strikingly, 5 of the 22 screened clones previously found to be noninfectious (30) were now infectious (S/N ≥ 5) and a further 8 of the 22 screened clones exhibited at least a five-fold increase in S/N (Table S1 and Fig. S2e). Finally, a comparison of calculated S/N for all screened clones from all three panels unequivocally demonstrated increased infectivity for HCVpp made in 293T^CD81KO^ cells (Fig. 2d & S2f). Combined, the above data suggest the 293T^CD81KO^ cells we have produced could be a useful resource for HCV research.

Having demonstrated improved infection, we sought to ensure deletion of CD81 in producer 293T cells did not affect the entry pathway of HCVpp. The E1 and E2 glycoproteins mediate the virus’ intricate entry pathway. The E1 and E2 genes display the greatest diversity in the HCV genome, yet receptor interactions remain highly conserved across distinct genotypes and entry is widely understood to involve virion interactions with the essential receptors CD81, SR-B1, Claudin and Occludin (35,36). We infected Huh-7 cells ablated for these receptors (37) with HCVpp produced in 293T^CD81KO^ cells. Whilst ablation of CD81, Claudin-1 and Occludin abolished infection for all four tested strains, a proportion of all strains was still able to infect SR-B1 knockout cells (Fig. S3). These data are consistent with previous findings for both HCVcc and HCVpp particles (37,38) and suggest pseudoparticles made in 293T^CD81KO^ cells follow the correct HCV entry pathway.

### Production in CD81 knock-out 293T cells does not alter HCVpp antibody sensitivity

HCVpp are widely used to examine the neutralizing breadth of and potency of monoclonal antibodies (mAbs) and polyclonal sera from humans and immunized animals (17,30). Therefore, it was imperative to ensure that HCVpp produced in 293T^CD81KO^ cells are antigenically similar to those produced in parental cells. First, we challenged H77 and UNKP 5.2.1 pseudoparticles with a serial dilution of AR3B, a bNAb that targets the CD81 binding site (CD81-bs) (39). We witnessed similar neutralisation by AR3B against both clones regardless of the cell line of production; indeed, the IC_50_ for the H77 particles was identical (Fig. 3a & 3b). We also challenged H77 and UNKP 5.2.1 pseudoparticles with a serial dilution of a soluble form of the large extracellular loop of CD81 (sCD81-LEL). As with AR3B, the IC_50_ of sCD81-LEL toward H77 was indistinguishable regardless of the cellular source of the HCVpp (Fig. 3c). The IC_50_ of sCD81-LEL toward the matched UNKP 5.2.1 HCVpp pair was also similar (Fig. 3d).

**Fig. 3.**
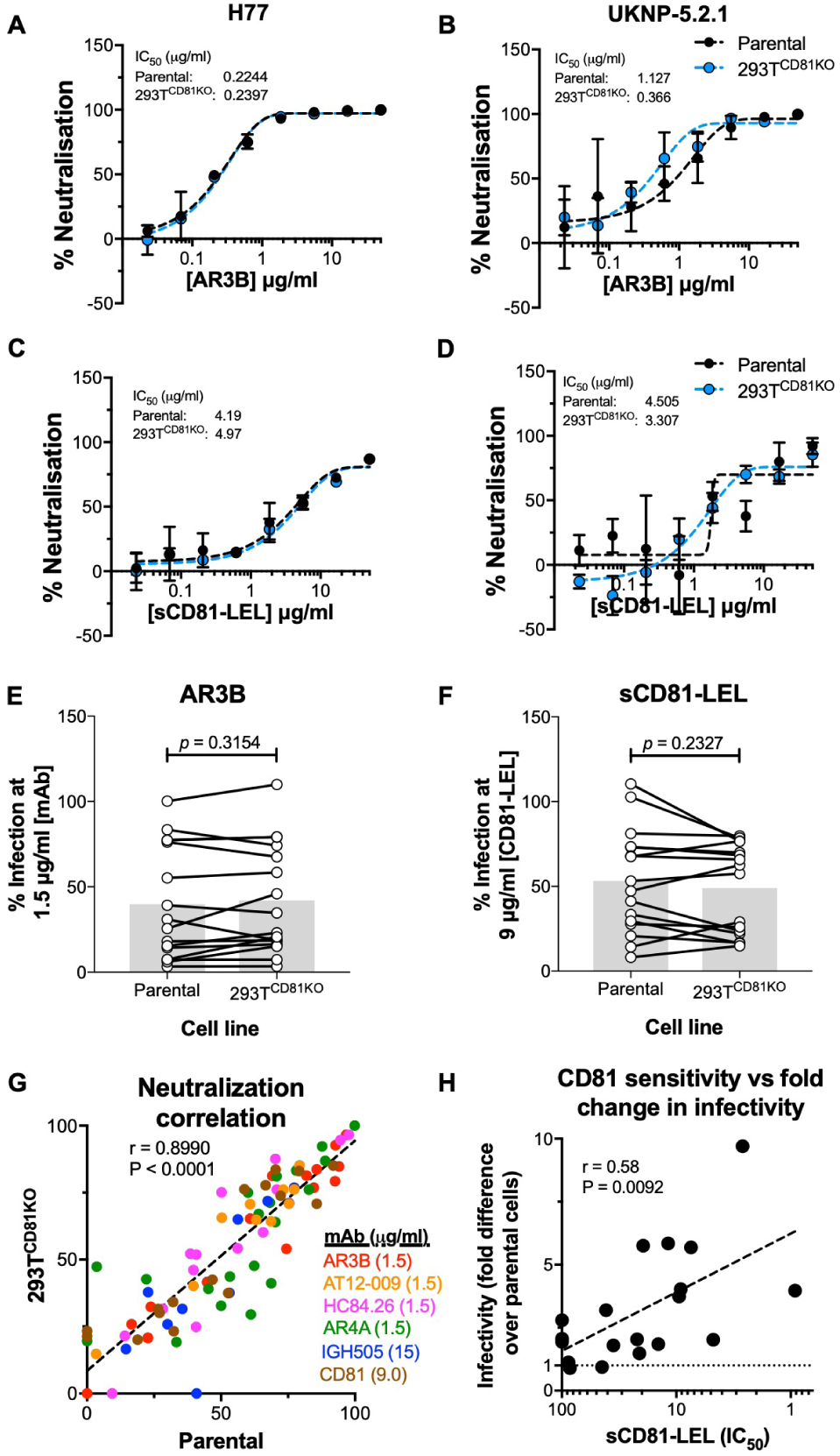
Absence of CD81 in producer 293T cells does not affect antibody sensitivity of HCVpp. The prototypical lab strain **(A)** and the clinical isolate **(B)**UKNP 5.2.1 were incubated with a serial dilution of the mAb AR3B prior to infection.The prototypical lab strain **(C)** H77 and the clinical isolate **(D)** UKNP 5.2.1 were incubated with a serial dilution of the sCD81-LEL. Data in a-d are expressed as percent neutralisation relative to wells not preincubated with antibody or receptor molecules and each point is the mean of three replicate values. Error bars indicate the standard deviations between replicate wells. Data were fitted using a sigmoidal curve in GraphPad Prism. A compiled summary of the infection levels of HCVpp panel consisting 16 clinical isolates preincubated with a single concentration of **(E)** AR3B (1.5 µg/ml) or **(F)** sCD81-LEL (9 µg/ml) prior to infection. Data are expressed as percent infection relative to wells not preincubated with antibody or receptor molecules. Connected points indicate a single E1E2 clone and the grey bars represent the mean value. Paired t-test, no significance. **(G)** A scatter plot of the percentage neutralisation for a panel of HCVpp clones produced in parental 293T cells (x-axis) vs when produced in 293T^CD81KO^ cells (y-axis) after challenge with a single concentration of indicated mAb or sCD81-LEL. **(H)** A scatter plot of HCVpp CD81 sensitivity (sCD81-LEL The IC_50_) (x-axis) and fold change in infectivity following production in 293T^CD81KO^ cells (y-axis). Each point in g and h represents a lone HCV isolate. Spearman correlation (r) and p values were computed on GraphPad Prism. All data are from lone experiments performed in triplicate.

Next, we compared the infectivity of an HCVpp panel following incubation with a fixed concentration of AR3B. We found the infectivity of most clones was similar irrespective of the cell line of production and saw no significant difference for the panel as a whole (Fig. 3e). This experiment was conducted for a further four mAbs; (i) AR4A, which targets an epitope composed of both E1 and E2 residues, (ii) AT12009 (CD81-bs), (iii) HC84.26 (domain D) and (iv) IgH505, which targets a discontinuous epitope in E1 (residues 313-327) (32,39–41). We saw no significant difference in the infectivities of the panel against all 4 mAbs irrespective of the cell line of production (Fig. S4). Similar observations were also made when pseudoparticles were pre-incubated with sCD81 (Fig. 3f). Finally, after generating a scatter plot for data shown in figures 3e, 3f and S4 we observed a very strong correlation for neutralisation sensitivity between HCVpp produced in 293T^CD81KO^ cells and HCVpp made in parental 293T cells. These data indicate ablation of CD81 in producer 293T cells does not change the sensitivity of pseudoparticles to neutralization by sCD81 or mAbs (Fig. 3g). However, we did find a significant positive correlation (spearman r = 0.58) between the sensitivity of HCVpp to sCD81 neutralization and their fold increase in infectivity following production in 293T^CD81KO^ cells (Fig 3h). This suggests that HCVpp that are sensitive to CD81 neutralization benefit more from production in 293T^CD81KO^ cells. In summary, the data demonstrate that producing HCVpp in 293T^CD81KO^ cells does not alter their antigenicity or sensitivity to mAbs. These findings considered along with their ability to improve virus infectivity suggest that the 293T^CD81KO^ cell line is a superior system for assessing the functionality and neutralisation of diverse HCVpp.

### sE2 produced in 293T^CD81KO^ or parental 293Ts displays a mixture of high-mannose and complex-type glycans

HEK 293T cells are commonly used for the production of sE2. Consequently, we examined whether 293T^CD81KO^ cells improve sE2 production as they did HCVpp production without altering the properties of the protein generated. To do this, we first adapted the 293T^CD81KO^ and parental 293T cell lines to grow in suspension, which makes the cells more suitable for large scale protein production. We refer to these cell lines as susp-293T and susp-293T^CD81KO^ cells. We used conventional transient transfection to produce sE2 in these two cell lines. In parallel, we also compared these to sE2 generated in the HEK293F cell lineage, which is a widely used cell line for producing recombinant proteins. We isolated sE2 monomers from harvested supernatants using affinity purification followed by size exclusion chromatography (SEC). Very similar yields of sE2 were obtained following purification of harvested susp-293T^CD81KO^, susp-293T (both ∼14 mg/L) and 293F (12 mg/L) supernatant, although relatively more aggregates were observed in sE2 from the susp-293T lineages. Furthermore, the retention volumes for monomeric sE2 made in susp-293T^CD81KO^ and susp-293T cells were practically identical (∼13.1 ml) yet monomeric sE2 produced in 293F cells eluted at a lower volume (∼12.3 ml) suggesting it is larger than that produced in the susp-293T cell lines (Fig. 4a). Indeed, SDS-PAGE analysis of unfractionated sE2 and their monomeric peak fractions, under both reducing and non-reducing conditions, revealed sE2 made in 293F cells migrated more slowly than sE2 from the two susp-293T lines (Fig. 4b). These data suggest that sE2 undergoes distinct post-translational modifications in 293F and 293T cells.

**Fig. 4.**
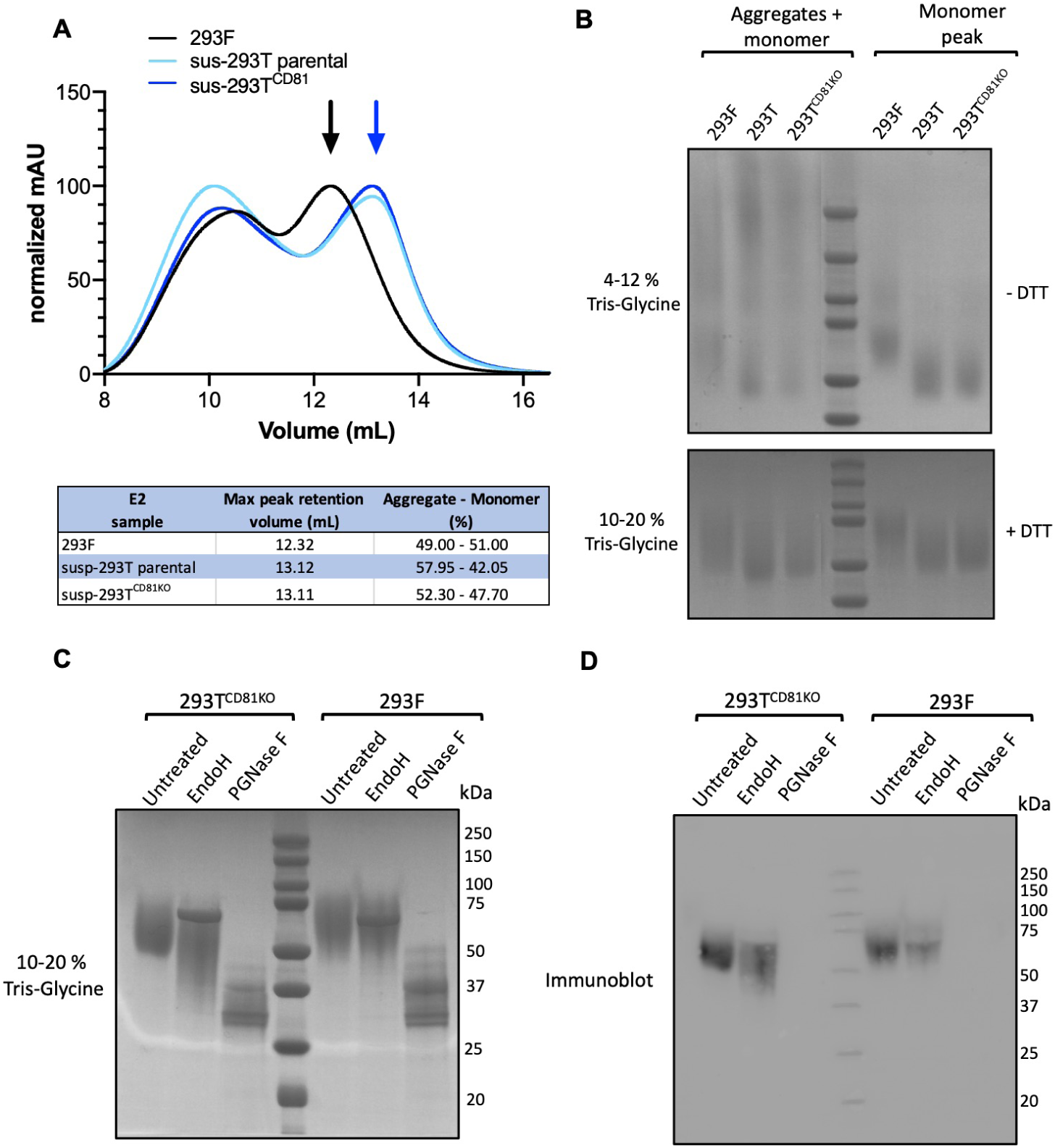
Soluble E2 is differentially glycosylated in susp-293T and 293F cells. **(A)** Superdex 200 size-exclusion chromatography profile of susp-293T(turquoise), susp-293T^CD81KO^ (blue) and 293F-(black) derived sE2 following Strep-II-tag affinity chromatography. Blue arrow corresponds to the peak fraction of monomeric sE2 from susp-293T and susp-293T^CD81KO^ cells and the black arrow corresponds to monomeric sE2 from 293F cells. mAU, milliabsorbance units. **(B)** Nonreducing BN-PAGE gel (top panel) and reducing SDS-PAGE gel (bottom panel) of the purified sE2 derived from the aforementioned cell lines (Coomassie Brilliant Blue G-250 staining). **(C)** Reducing SDS-PAGE gel of monomeric susp-293T^CD81KO^- and 293F-derived sE2 after treatment with PGNase F or EndoH (Coomassie Brilliant Blue G-250 staining). **(D)** Western blot of SDS-PAGE gel shown in C. Molecular mass markers are indicated on the right (kDa). Data is from a single experiment.

Up to 11 Asn-linked glycosylation sites can be detected in most E2 sequences and fully processed E2, in both its soluble form or in the context of viral particles, is heavily glycosylated and displays a mixture of high-mannose and complex-type glycans (42,43). As a result, the molecular mass of E2 is significantly influenced by glycans. To test whether differences in glycosylation account for the disparate molecular weight between sE2 from 239T and 293F cells, we treated purified monomeric sE2 with PGNase F or EndoH deglycosylation enzymes. PGNase F cleaves both high-mannose and complex-type glycans yielding a mostly deglycosylated protein whereas Endo H specifically cleaves high-mannose and some hybrid glycans.

We compared sE2 produced in 293F to susp-293T^CD81KO^-derived sE2 (glycoprotein produced in either the edited or parental 293T cells migrated identically suggesting their glycosylation status is similar, Fig. 4b). As expected, PGNase F treatment confirmed sE2 made in both susp-293T^CD81KO^ cells and 293F cells to be heavily glycosylated. Undigested sE2 migrated to around ∼65 kDa and its deglycosylated form migrated to ∼35 kDa irrespective of the cell line of production (Fig. 4c). Strikingly, the sensitivity of sE2 from 293F and susp-293T^CD81KO^ cells to EndoH treatment was different. sE2 made in 293F cells was mostly unaffected by EndoH treatment as evidenced by the similar migration observed for the untreated and EndoH digested samples (Fig. 4c). This suggests that 293F-produced sE2 is composed of mostly complex-type glycans, a hallmark of maturation through the Golgi apparatus. On the other hand, sE2 from susp-293T^CD81KO^ was sensitive to EndoH as evidenced by the faster migrating smear on SDS-PAGE after treatment. Furthermore, the smear length indicates glycans on 293T-produced sE2 ranged from fully EndoH resistant to almost completely EndoH sensitive. This suggests that susp-293T^CD81KO^ cells produce a heterogeneous population of sE2 molecules in terms of glycosylation and that some sE2 molecules are only partially matured as some remain high-mannose type, while others have been heavily modified by Golgi enzymes (Fig. 4c & 4d). Notably, heterogeneity in the glycosylation status of E2 has also been observed in HCVcc-associated E2 (42). This implies that the processing of sE2 through the Golgi apparatus of susp-293T^CD81KO^ cells more closely resembles that of E2 destined to be incorporated in a full-length virion than sE2 from 293F cells. In summary, these data demonstrate compositional differences in the glycans of sE2 produced in susp-293T^CD81KO^ and 293F, based on previous findings it is likely that susp-293T^CD81KO^-produced sE2 is more representative of E2 on HCVcc particles.

### CD81 binding and antigenicity of 293T lineage produced sE2 is superior to that of 293F-produced sE2

To investigate whether producing sE2 in cells lacking CD81 altered the glycoprotein epitope presentation, we first tracked the binding of immobilised sE2 monomers from susp-239T, susp-293T^CD81KO^ and 293F cells to serially diluted mAbs and human Fc-tagged soluble CD81 (sCD81-LEL-hFc) by enzyme-linked immunosorbent assay (ELISA). The mAb panel included the AS412 targeting AP-33, AR3B and AT12-009 which target the front layer and binding loop of the CD81 binding site, CBH-4B (domain A) and HC84.26 (domain E) (32,39,40,44). All tested antibodies and sCD81-LEL-hFc bound equally well to sE2 produced in susp-293T^CD81KO^ or parental susp-293T cells, suggesting that the expression of CD81 in 293T cells does not affect the antigenicity of sE2 (Fig. 5a). sE2 produced in 293F cells exhibited reduced binding to conformational antibodies, with equivalent binding observed only for AP33, which targets a continuous epitope (AS412). These results suggest there is a difference in epitope accessibility between the two forms of sE2. This difference was even more apparent when comparing the binding of sCD81-LEL-hFc to the different sE2 types (Fig. 5b).

**Fig. 5.**
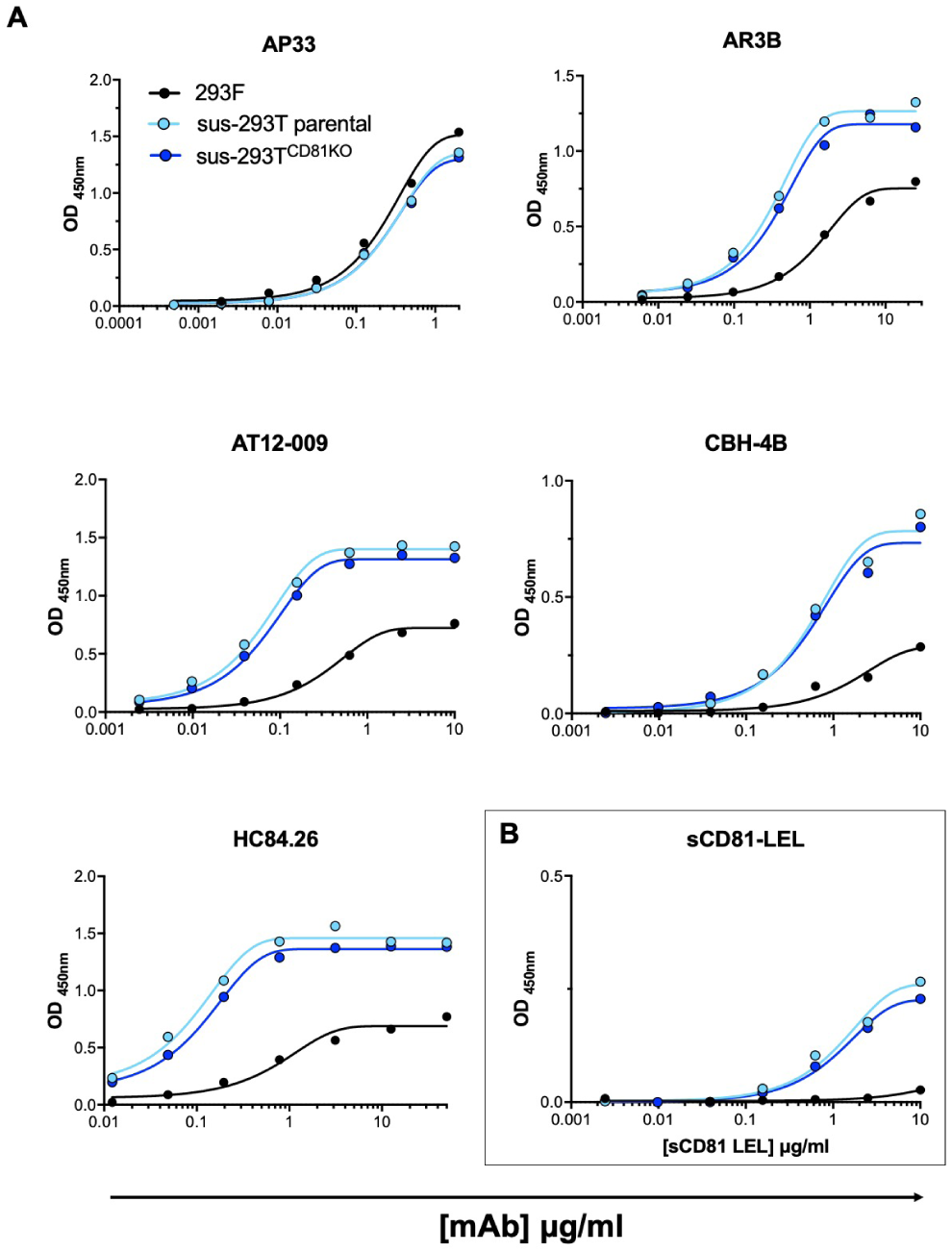
ELISA analysis of antibody and CD81 binding to sE2 form susp-293T, susp-293T^CD81KO^ and 293F cells. sE2 were immobilised to Strep-TactinXT coated microplates and incubated with a serial dilution of the indicated mAb (a) or CD81-LEL-hFc (b). Each point represents a single value from a lone experiment. Each point represents a single value from a lone experiment.

The binding of the three types of sE2 to the mAb panel and sCD81-LEL-hFc was further characterized by biolayer interferometry (BLI) (Fig. 6). Antibodies and CD81-LEL-hFc were immobilized on a protein A biosensor and incubated with the same concentration of sE2. Consistent with the ELISA data, near-identical binding profiles were observed for sE2 made in susp-293T^CD81KO^ and parental susp-293T cells; again, besides AP33, sE2 made in 293F exhibited lower binding to all mAbs and sCD81-LEL. Together, the BLI and ELISA data reveal a previously unappreciated difference between the antigenicity of sE2 produced in 293F and 293T cells. This may have implications for functional/structural studies of sE2 and/or future immunogen production.

**Fig. 6.**
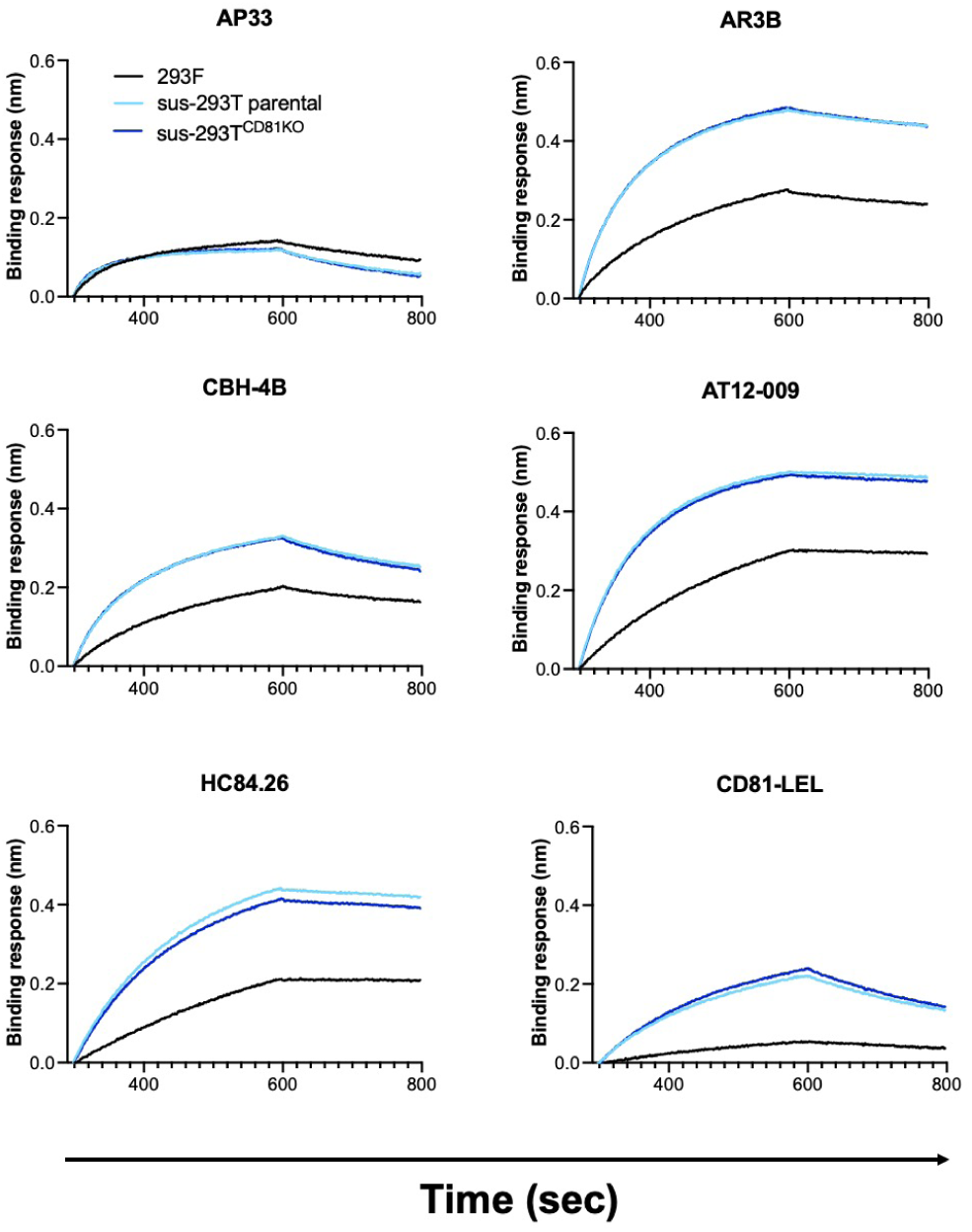
BLI analysis of antibody and CD81 binding to sE2 form susp-293T, 293T^CD81KO^ and 293F cells. CD81-LEL-hFc and the indicated mAbs were immobilised onto protein A biosensors and incubated with a single concentration of sE2. Sensorgrams were obtained from an Octet K2 instrument. Data is from a single experiment.

## Discussion

The HCV pseudoparticle system is a flexible platform for evaluating the phenotypes of clinical isolates and has proven to be a crucial tool for defining the sequence of virion and receptor interactions occurring during HCV entry; it is likely to be an important component for the assessment of future HCV vaccine candidates (13,28,29). Nonetheless, producing infectious viruses from polymorphic HCV populations circulating in patients has repeatedly proved challenging (33,45,46). This is problematic for vaccine screens and antibody neutralizing breadth studies, as only strains which recover workable infectivity are included in such analyses and they may not necessarily be representative of the diversity circulating in the population. Moreover, some infectious strains give low yields which fail to reach a signal to noise threshold sufficient for neutralisation analysis, a signal at least 10-fold above background is favoured but can be lowered to a minimum of 5 for clinical isolates (30,47). Perhaps owing to its flexibility, several studies have demonstrated the HCVpp system is amenable to optimisation. Urbanowicz, Tarr and colleagues have shown infectivity for some strains in the HCVpp system partly depends on the choice of retroviral background and the stoichiometry of transfected plasmids (31). More recently, it has been shown that the cellular co-expression of the HCV p7NS2 open reading frame in producer cells enhances pseudoparticle infectivity for the H77 strain (48). Others have also proposed that the inclusion of an intact HCV Core at the N-terminal of the glycoprotein plasmid is required for efficient pseudoparticle morphogenesis for some strains (49). However it is now generally recognised that the inclusion of the last 21 residues of the Core/E1 peptide is sufficient for the vast majority of clones. Here, we disclose that a 293T cell lineage genetically edited to not express CD81 improves or rescues the infectivity of a wide range of HCVpp strains.

Producing HCVpp in 293T^CD81KO^ cells is a facile method for improving HCVpp infectivity as it does not require empirical optimization of the plasmid ratios or co-expression of additional HCV genome elements. Unlike Soares and colleagues, we demonstrate a blanket increase for the infectivities of a broad range of strains representing multiple genotypes (48). Furthermore, producing HCVpp in 293T^CD81KO^ cells had a more dramatic effect on viral infectivity compared to plasmid ratio optimization (31). Moreover, none of the 45 clones showed a decrease in infectivity after production in 293T^CD81KO^ (see Table S1). Therefore we would recommend the routine adoption of this cell system for the generation of HCVpp and are happy to provide this cell line to any interested investigators.

We are yet to uncover the mechanism by which the presence of CD81 in the 293T cells influences the infectivity of pseudoparticles. However, the correlation between HCVpp sCD81 neutralization sensitivity and the fold increase in infectivity following production 293T^CD81KO^ cells (Fig. 3H) may suggest that there is a direct interaction between CD81 produced by regular 293T cells and HCVpp. There are several points during HCVpp production where E2 could encounter its cognate binding partner. The E1E2 glycoprotein and CD81 undergo glycosylation and palmitoylation in the Golgi compartment, respectively. When co-transfected, CD81 associates with E1E2 glycoprotein in the endoplasmic reticulum (ER) and influences the glycoprotein’s maturation through the Golgi membranes (50). This association redirects the glycoprotein toward the endocytic pathway culminating in its incorporation into exosomes possibly reducing the amount of E1E2 available at the cell surface, where retroviral budding occurs. CD81 is also incorporated into retroviral pseudoparticles and its deletion possibly reduces protein density in lipid rafts at the cell membrane thereby permitting for increased E1E2 incorporation into budding HCVpp. However, glycoprotein incorporation is a poor predictor of HCVpp infectivity as others have shown that the majority of E2 in the supernatant of cells producing pseudoparticles does not sediment with infectious particles (51). Additionally, as released virions in suspension will behave as colloidal matter (52) it is highly probable that many particles contact the cell surface and may be retained or fated to non-productive uptake since 293T cells do not express the additional HCV entry factor claudin (28,29). In preliminary experiments, we have observed that parental 293T conditioned media, which contains extracellular CD81, is detrimental to the infectivity of HCVpp produced in 293T^CD81KO^ cells and not to HCVpp produced in parental 293Ts (data not shown). This support a cell-free interaction between HCVpp and CD81, possibly in the context of exosomes.

Our findings somewhat contrast a recent report by Soares and colleagues who concluded that CD81 was required to generate HCVpp (53). This is likely because Soares et al. did not perform infections with the HCVpp they generated from cells silenced for CD81 expression and instead relied on the quantities of E2 in viral harvests as their gauge. Furthermore, as mentioned above, E2 incorporation is a poor predictor of HCVpp infectivity (51) and the fall in E2 quantities they observed in their HCVpp harvests following CD81 silencing is more likely a reflection of reduced E2 incorporation into exosomes since CD81 chaperons E2 into the endocytic pathway, from where exosomes are biosynthesized (50).

Considering CD81 ablation evidently improves the production of HCVpp, it is surprising we did not observe an increase in the amount sE2 recovered from susp-293T^CD81KO^. This may be a consequence of the absence of the signal peptide at the intersection of core and E1, which retains E1E2 in the ER lumen, where virion morphogenesis would normally commence in full-length virus (54). Therefore it is plausible that, unlike the E1E2 heterodimer, sE2 has reduced opportunity for interaction with CD81 as it matures through the Golgi membranes.

The observation that sE2 made in susp-293T and 293F cells is differentially glycosylated was unexpected and requires further scrutiny. EndoH digestion indicated some E2 glycans remain shielded from Golgi glycosidases and glycosyltransferases in 293Ts, whereas a much higher proportion of total sE2 efficiently matured through the Golgi apparatus when produced in 293Fs. Differential glycosylation patterns for the same viral glycoprotein have been demonstrated in cell lines from different species but not in cell lines sharing an immediate predecessor, as 293Ts and 293Fs do (43,55,56). There are likely many differences between the two lines; however, the most obvious are that 293Fs are usually grown in suspension and do not express Simian virus large T 40 antigen. We adapted the 293T^CD81KO^ to suspension thus this cannot explain the difference in glycosylation. The SV40 origin of replication is present on the H77 sE2 expression vector (pPPI4) we used (57); however, it is highly improbable that an improvement in plasmid stability would affect the distal process of sE2 glycosylation. A recent omics study of HEK293 and its progeny cell lines not only revealed transcriptome profile differences between adherent 293T cells and 293F cells but also between 293H cells adapted to suspension and 293F cells (58). This indicates genes other than those involved in the adherent to suspension transition are also differentially expressed 293 progeny cell lines. Hence we suspect some genetic difference(s) between 293F and 293T cells ultimately underlies our observation.

The observed disparity in the antigenicity between 293T- and 293F-produced sE2 is probably a direct result of the difference in glycosylation pattern between the two. Glycans, directly and indirectly, influence E1E2 folding through their interactions with ER chaperones such as calnexin (59,60). Several E2 glycans have been shown to play an important role in folding and E1 and E2 heterodimerization (61). Furthermore, it is well documented that glycans protect conserved viral glycoprotein epitopes, including those on E1E2, from recognition by neutralizing antibodies through a phenomenon termed glycan shielding (60,62,63). Our deglycosylation experiments indicate that 293F-produced sE2 molecules harbour mostly complex glycans, while susp-293T^CD81KO^-produced sE2 also contains high-mannose or hybrid glycans. Complex glycans are usually larger and this could explain why some epitopes on 293F-produced E2 seem to be less accessible for most mAbs and CD81 as measured by ELISA and BLI (Fig. 5 and Fig. 6). Our results suggest that apparently highly similar 293-derived cell lines can produce glycoproteins with different glycan species. Sitespecific glycan analysis (43,64) is needed to more thoroughly compare the difference in glycosylation between two lines and its effect on antigenicity.

Deleting one of the main receptors of HCV from producer cells significantly increased HCVpp infectivity. It is plausible that similar receptor-deleting strategies could be applied to increase the recovered infectious titres of other pseudotyped viruses. However, one should always carefully consider the biological role of these receptors. For example, integrins and lipoprotein receptors are targeted by a range of viruses, but integrins also play a significant role in maintaining cell integrity and cell cycle progression, while many lipoprotein receptors are essential for proper cholesterol homeostasis (65,66). Disruption of either cellular processes would likely reduce infectious titres. Therefore, yielded viruses must be phenotyped for cell entry pathway and antibody sensitivity to ensure there are no disparities with pseudoparticles obtained from the parental lineage, as we have done here.

In summary, here we have demonstrated that the presence of the HCV receptor CD81 in producer 293T cells used for HCVpp production can affect recoverable titres. We proceeded to generate a new 293T cell line lacking CD81 that substantially improved infectivity for most clinical HCVpp isolates. Crucially, these HCVpp are functionally similar to those made in conventional 293Ts. This cell line can easily be generated through CRISPR-Cas9 gene-editing but is also readily available from us. Furthermore, we disclose differences in the glycosylation patterns and antigenicity of sE2 produced in 293F and 293T cells; this could have important implications for functional studies of E2 and for immunogen production.

## Supporting information

Kalemera M et al 2020 suppl. info

## Materials and methods

### Cell lines and culture

The receptor knock-out Huh-7 celllines were a kind gift from Prof. Yoshiharu Matsuura (Osaka University). Huh-7 cells were acquired from Apath LLC. All cells were grown at 37^*°*^ C in Dulbecco’s Modified Eagle Medium (DMEM) (Gibco) supplemented with 10% foetal calf serum (FCS), 1% non-essential amino acids and 1% penicillin/ streptomycin (P/S) (DMEM/FCS). For soluble E2 production, HEK293T (ATCC) and 293T^CD81KO^ cells were initially maintained in DMEM/FCS. For large-scale purifications, these cells were adapted to suspension culture by growing the cells in a mix of FreeStyle(tm) 293 Expression medium (Gibco) and DMEM/FCS and decreasing the concentration of DMEM/FCS at each passage. Finally, both cell lines were grown in pure Freestyle medium. HEK293F cells (Invitrogen) were maintained in FreeStyle(tm) 293 Expression. All suspension cell lines were maintained in a shaking incubator (125 rpm) at 37^*°*^ C and 8% CO_2_.

### Antibody source table

Antibody source table.
| Antibody | Source | DOI |
| --- | --- | --- |
| Goat anti-human IgG (HRP) | Jackson Immunoresearch | N/A |
| Goat anti-mouse IgG (HRP) | CST | N/A |
| 2.131 (anti-CD81) | Grove et al., 2017 (67) | 10.12688/wellcomeopenres.12058.1 |
| AP33 | Clayton et al., 2002 (68) | 10.1128/JVI.76.15.7672-7682.2002 |
| AR3B | Law et al., 2008 (69) | 10.1038/nm1698 |
| AR4A | Giang et al., 2012 (39) | 10.1073/pnas.1114927109 |
| AT12-009 | Merat et al., 2016 (32) | 10.1371/journal.pone.0165047 |
| CBH-4B | Keck et al., 2004 (70) | 10.1128/JVI.78.17.9224-9232.2004 |
| H77.16 | Sabo et al., 2011 (71) | 10.1128/JVI.00586-11 |
| HC84.26 | Keck et al., 2012 (40) | 10.1371/journal.ppat.1002653 |
| IGH505 | Meunier et al., 2008 (41) | 10.1128/JVI.01872-07 |

### Western blot

Two days prior to study, HEK 293T and Huh-7 cells +/-CD81 were seeded into a standard 6 well plates at 2.5×10^5^ cells/ml. Cells were then lysed using a buffer containing 20mM Tris-HCl, 135mM NaCl, 1% Triton-X 100 and 10% glycerol. The samples were then run on a TruPage 4-12% gel under non-reducing conditions and transferred on to nitrocellulose membrane. The blots were blocked in PBS + 2% milk solution + 0.1% Tween-20 and then probed by serial incubation with anti-receptor antibodies and goat anti-mouse secondary conjugated to horseradish peroxidase (HRP). Chemiluminescence signal was then measured in a Chemidoc MP (Bio Rad).

### Generation of CD81 knock-out cell lines

293T^CD81KO^ cell lines were established by following the Invitrogen TrueGuide(tm) sgRNA CRISPRMAX(tm) transfection protocol. Briefly, 293T cells were first transfected with one of the following CD81 sgRNAs: CRISPR661076_CR, CRISPR661085_CR and CRISPR661093_CR (Thermo Fisher Scientific). Seventy-two hours later, the cells were then sorted on a FACSAria(tm) II instrument (BD Biosciences) to obtain CD81-/-populations. Cells were then expanded for 48 hours before dilution for single cell cloning and expansion in a 96 well plate. Knock-out of CD81 was confirmed by flow cytometry and Western blot.

### Generation of HCV pseudoparticles

To generate HCVpp, HEK293T cells were co-transfected with three plasmids: an HIV (pCMV-dR8.91) or MLV (phCMV-5349) packaging construct, a luciferase reporter plasmid and an expression vector encoding the appropriate HCV glycoprotein. A no-envelope control (empty plasmid) was used as a negative control in all experiments. Supernatants containing HCVpp were collected at 48 and 72 hours.

### Infection and Neutralisation assays

Huh-7 cells were seeded at 1.5 × 10^4^ cells per well of a 96 well plate 24 hours prior to the experiment; to infect cells were challenged with HCVpp supernatant. For neutralization studies, a volume of virus stock was incubated in triplicates with a dilution series of anti-E2 antibody or sCD81-LEL prior to infection. Virus-antibody or virus-sCD81-LEL mixtures were incubated for 1 hour at 37^*°*^ C and were added to Huh-7 cells and infection was allowed to proceed for 72 hours. For readout, the samples were lysed and assayed using the SteadyGlo reagent kit and a GloMax luminometer (Promega, USA). A no-envelope pseudovirus was used to calculate the signal-to-noise ratio. Data was analysed in GraphPad Prism 7.0c.

### Soluble E2 protein expression and purification

Strep-II-tagged soluble E2 (sE2) was expressed using the same transient transfection protocol in all three HEK293-based cell lines. The supernatant containing sE2 was harvested after six days and clarified using vacuum-filtration (0.2 *µ* m) and sE2 was purified using Strep-TactinXT columns (IBA Life Sciences) by gravity flow (∼0.5-1.0 ml supernatant/min). The eluted proteins were then concentrated and buffer exchanged into Tris-buffered saline (TBS, pH 7.5) using Vivaspin 6, 10.000 MWCO PES (Sartorius). Finally, purified sE2 was fractionated using size-exclusion chromatography (SEC) (Superdex 200 Increase 10/300GL, GE) and the fractions corresponding to E2 monomers were pooled and concentrated. The concentration of the monomer fraction was then determined by Nanodrop (Thermo Fisher Scientific) using theoretical molecular weight and extinction coefficient.

### Blue native (BN-)PAGE and SDS-PAGE

BN-PAGE and SDS-PAGE analyses were performed as described elsewhere (72) with some modifications. Briefly, for BN-PAGE, 5 *µ*g sE2 was mixed with PBS and 4x loading dye (a mix of 500 *µ*l 20x MOPS Buffer (1M MOPS + 1M Tris, pH 7.7), 1000 *µ*l 100% glycerol, 50 *µ*l 5% Coomassie Brilliant Blue G-250, 600 *µ*l milli-Q) and directly loaded onto a 4-12% BisTris NuPAGE gel (Thermo Fisher Scientific). The gels were run for 1 hour at 200 V at 4C using NativePAGE Running Buffer (Invitrogen). BN-PAGE gels were stained using the Colloidal Blue Staining Kit according to the manufacturer’s instructions (Life Technologies). For SDS PAGE, 5 *µ*g of sE2 was mixed with loading dye (25 mM Tris, 192 mM Glycine, 20% v/v glycerol, 4% m/v SDS, 0.1% v/v bromophenol blue in milli-Q water) and incubated at 95^*°*^ C for 10 min prior to loading on a 4-12% Tris-Glycine gel (Invitrogen). For reducing SDS-PAGE, dithiothreitol (DTT; 100 mM) was included in the loading dye and loaded on a Novex 10-20% Tris-Glycine gel (Thermo Fisher Scientific). Gels were run in a buffer containing 25 mM Tris, 192 mM glycine and 0.5% SDS for 1.0h at 200 V at 4^*°*^ C. Coomassie blue staining of SDS-PAGE gels was performed using the PageBlue Protein Staining Solution (Thermo Fisher Scientific).

### Deglycosylation of purified E2 monomers

Purified sE2 monomers were treated with PNGase F or EndoH (New England BioLabs) following the manufacturer’s protocol. The samples were then run on a reducing Novex 10-20% Tris-Glycine gel (Thermo Fisher Scientific) and stained by Coomassie blue as described above. Hereafter, the samples from this same gel were transferred onto a nitrocellulose membrane. The blots were blocked in PBS + 5% milk solution + 0.1% Tween-20 and then probed by serial incubation with anti-E2 antibody and goat anti-mouse secondary conjugated to HRP. Chemiluminescence signal was then determined via the use of the Western Lightning Plus-ECL system (PerkinElmer).

### Enzyme-linked immunosorbent assay (ELISA)

Purified StrepII-tagged sE2 monomer (1.0 *µ*g/mL in TBS) was coated for 2 hour at room temperature on 96-well Strep-TactinXT coated microplates (IBA LifeSciences). Plates were washed with TBS twice before incubating with serially diluted mAbs and CD81-LEL-hFc (R& D systems) in casein blocking buffer (Thermo Fisher Scientific) for 90 min. After three washes with TBS, a 1:3000 dilution of HRP-labelled goat anti-human IgG (Jackson Immunoresearch) in casein blocking buffer was added for 45 min. After washing the plates five times with TBS+0.05% Tween-20, plates were developed by adding develop solution (1% 3,3’,5,5’-tetraethylbenzidine (Sigma-Aldrich), 0.01% H_2_O_2_, 100 mM sodium acetate, 100 mM citric acid) and the reaction was stopped after 3 min by adding 0.8 M H_2_SO_4_. Absorbance was measured at 405/570nm and data was visualized and analysed on GraphPad Prism.

### Biolayer interferometry (BLI)

BLI assays were performed using the Octet K2 instrument (FortéBio). All assays were performed at 30^*°*^ C and with agitation set at 1000 rpm. Antibodies, CD81-LEL-hFc and sE2 samples were dissolved in running buffer (phosphate-buffered saline (PBS) pH 7.5 with 0.1% bovine serum albumin (BSA) and 0.02% Tween-20 in a volume of 250 *µ*l/well. Antibodies (3.0 *µ*g/mL) and CD81-LEL-hFc (5.0 *µ*g/mL) were immobilized onto protein A biosensors (FortéBio, cat no. 18-5010) until a loading threshold of 1.0 nm was reached followed by a 30 sec baseline measurement in the running buffer. Purified sE2 monomers were diluted to 250 nM and association and dissociation were measured for 300 sec and 200 sec, respectively. A well without E2 was used for background correction. Data was analysed and visualized in GraphPad Prism.

### Statistical analysis

All statistical analysis was performed on GraphPad Prism.

## Author contributions

*Conceptualization*: MDK, KS, JG. *Formal analysis*: MDK, JCP, AC, AU. *Funding acquisition*: RAB, JS, JG. *Investigation*: MDK, JCP, AC, AU. *Methodology*: RAB, KS, JG. *Project administration*: MDK, RAB, KS, JG. *Resources*: RAB, JS, JG. *Supervision*: RAB, KS, JG. *Validation*: RAB, KS, JG. *Visualization*: MDK, JCP, AC, AU. *Writing – original draft*: MDK. *Writing – review* & *editing*: All authors.

## Acknowledgements

We are grateful to Alexander Tarr and Jonathan Ball for providing E1E2 expression plasmids for HCVpp with the prefix UKNP. We also thank the Fondation Dormeur, Vaduz and Marit J van Gils for funding and acquiring the Octet K2 instrument.

