## Supplementary material for "Optimised cell systems for the investigation of hepatitis C virus E1E2 glycoproteins": Kalemera M et al 2020 suppl. info

Kalemera, Capella-Pujol, Chumbe et. al.

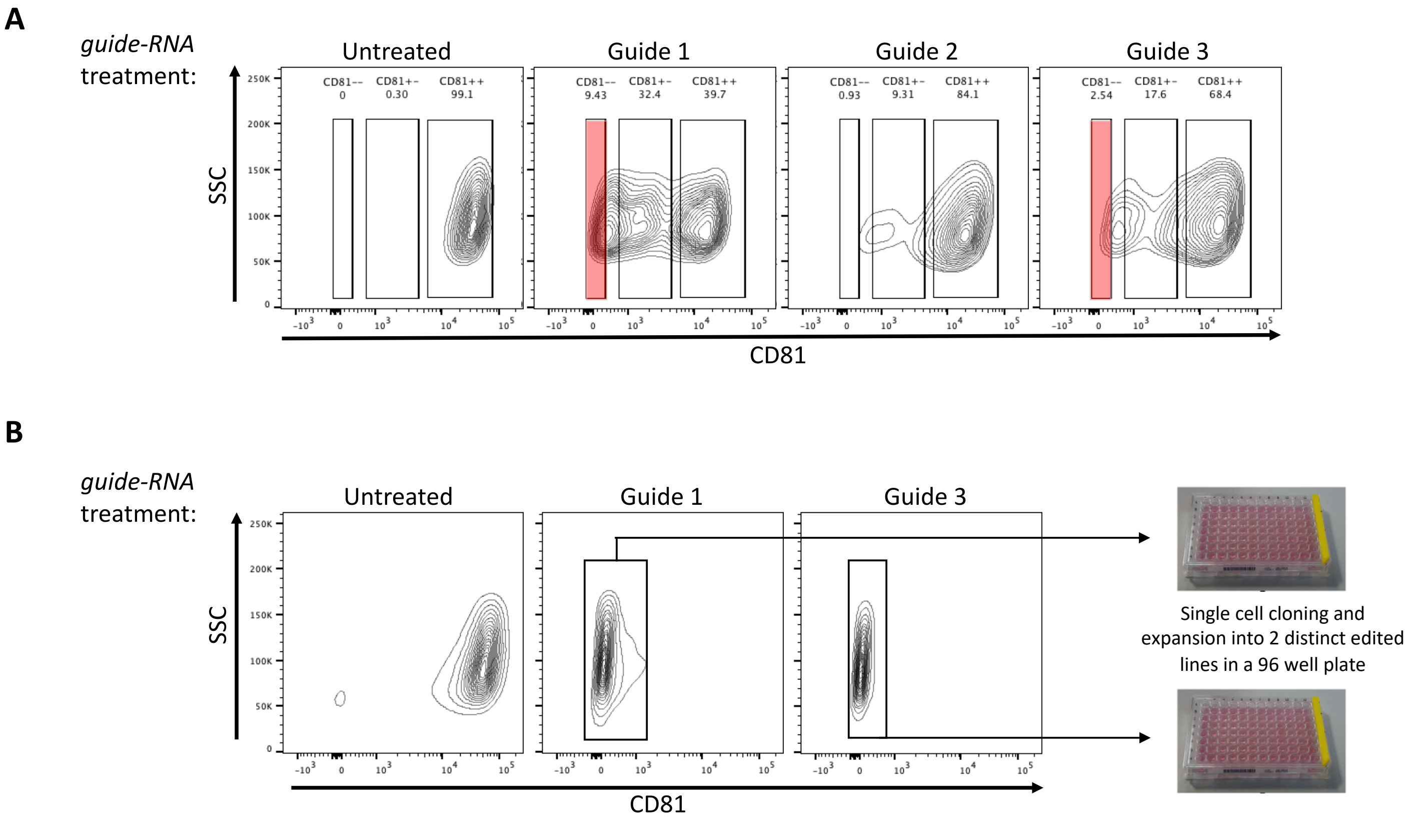


**Supplementary Figure 1. Generation of CD81 knock-out 293T cell lines.** 293T cells were transfected with CRISPR Cas9 gene-editing components along with one of three guide RNAs (sgRNA) targeting the CD81 exon. **(A)** 72 hours later cells were sorted using flow cytometry to obtain CD81 double negative populations (CD81-/-) (red coloured boxes). **(B)** Cells that received sgRNAs guide 1 and 3 were then expanded for 48 hours before dilution for single cell cloning and expansion in a 96 well plate.

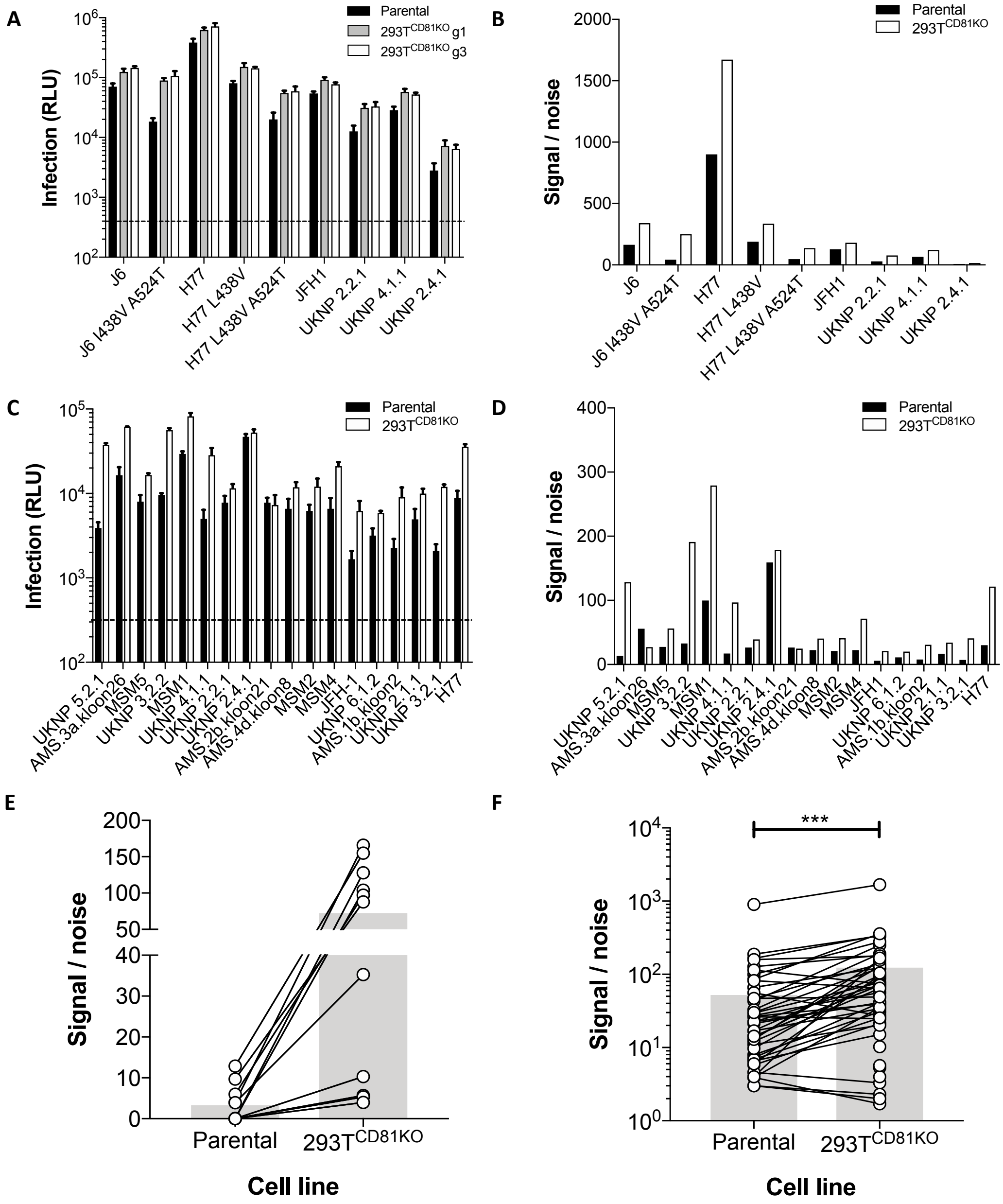


**Supplementary Figure 2. HCVpp made in CD81 knock-out 293T cells exhibit enhanced infectivity.** Huh-7 cells were challenged with HCVpp produced in parental or $293T^{CD81KO}$ cell lines. The infection levels **(A)** and calculated S/N **(B)** for an HCVpp panel consisting of both prototypical and clinical isolates. Note, only S/N of virus made in $293T^{CD81KO}$ g3 cells is included in b. The infection levels **(C)** and calculated S/N **(D)** of a clinical isolate HCVpp panel. Error bars in a and c indicate the standard deviations between three replicate wells. **(E)** Stratification of the top improving clones and clones whose infection was rescued when produced in $293T^{CD81KO}$ cells. **(F)** A compiled summary of the calculated S/N for all screened clones (see Table S1). Paired t-test, (***$p < 0.001$). Connected points indicate a single E1E2 clone and the grey bars represent the mean value. All data are from lone experiments performed in triplicate. Dash line indicates background signal determined by infection of pseudoparticle lacking E1E2.

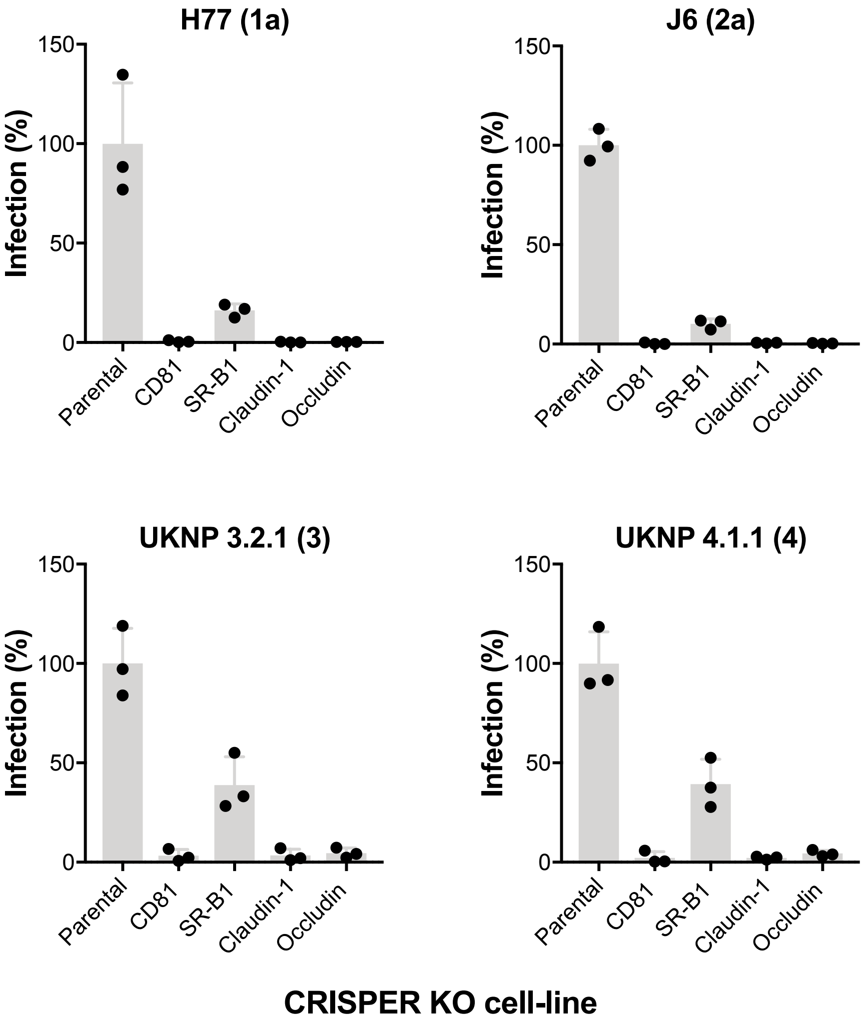


**Supplementary Figure 3. Receptor dependency of 293T$^{CD81KO}$ produced HCVpp**. Huh-7 cell lines CRISPR/Cas9 engineered to not express one HCV receptor were challenged with pseudoparticles carrying E1E2 glycoproteins from HCV genotypes one to four. Data are expressed as a percentage of the infection observed in parental Huh-7 cells. Error bars indicate the standard deviations between replicate wells. Data is from a single experiment performed in triplicate.

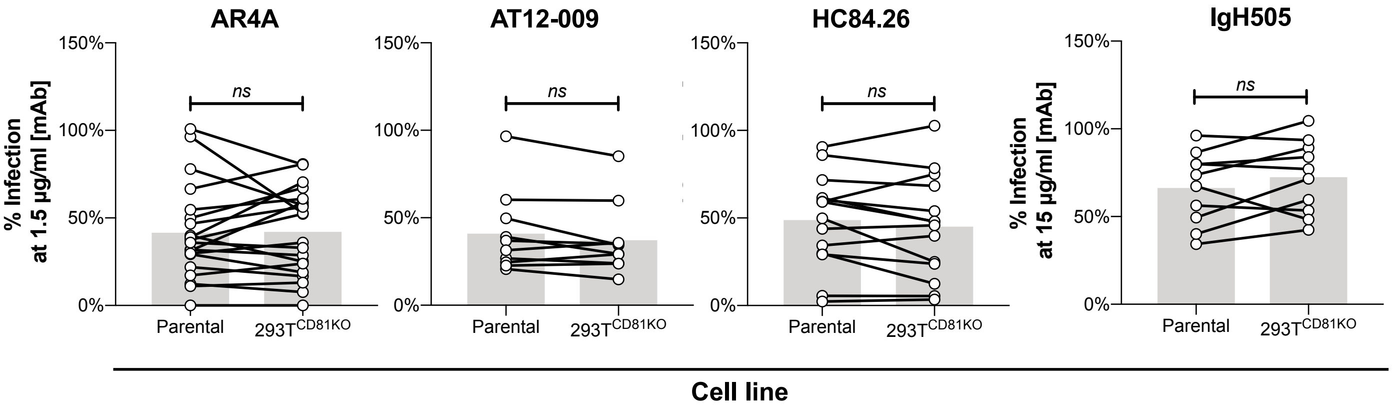


**Supplementary Figure 4. Absence of CD81 in producer 293T cells does not affect antibody sensitivity of HCVpp.** Compiled summaries of the infection levels of HCVpp when preincubated with a single concentration of the indicated mAb prior to infection. Connected points indicate a single E1E2 clone and the grey bars represent the mean value. Paired t-test, no significance (ns). All data are from lone experiments performed in triplicate.

| HCVpp [a] | Core [b] | Genotype | 293T line of HCVpp production | | Fold change in S/N [c] |
|---|---|---|---|---|---|
| | | | Parental | 293T$^{CD81KO}$ | |
| H77 | Lentiviral | 1a | 901.34 | 1673.03 | 1.86 |
| H77 L438V ✤ | Lentiviral | 1a | 188.55 | 335.28 | 1.78 |
| H77 L438V A524T ✤ | Lentiviral | 1a | 47.04 | 138.06 | 2.94 |
| J6 | Lentiviral | 2a | 164.88 | 340.21 | 2.06 |
| J6 I438V A524T ◆ | Lentiviral | 2a | 43.05 | 250.15 | 5.81 |
| JFH1 | Lentiviral | 2a | 127.48 | 181.22 | 1.42 |
| UKNP 2.2.1 | Lentiviral | 2 | 29.57 | 77.21 | 2.61 |
| UKNP 2.4.1 | Lentiviral | 2 | 6.57 | 15.02 | 2.29 |
| UKNP 4.1.1 | Lentiviral | 4 | 66.53 | 122.53 | 1.84 |
| AMS.1b.kloon2 | Retroviral | 1b | 7.96 | 30.86 | 3.88 |
| AMS.2b.kloon21 | Retroviral | 2b | 26.45 | 24.91 | 0.94 |
| AMS.3a.kloon26 | Retroviral | 3a | 55.90 | 27.27 | 0.49 |
| AMS.4d.kloon8 | Retroviral | 4d | 22.47 | 40.43 | 1.80 |
| h77 | Retroviral | 1a | 30.11 | 121.48 | 4.03 |
| jfh1 | Retroviral | 2a | 5.79 | 21.29 | 3.68 |
| msm1 | Retroviral | 1 | 99.77 | 279.03 | 2.80 |
| msm2 | Retroviral | 2 | 21.22 | 41.13 | 1.94 |
| msm4 | Retroviral | 4 | 22.46 | 71.53 | 3.19 |
| msm5 | Retroviral | 5 | 27.34 | 56.30 | 2.06 |
| uknp 2.1.1 | Retroviral | 3 | 26.61 | 39.23 | 1.47 |
| uknp 2.2.1 | Retroviral | 2 | 159.01 | 178.74 | 1.12 |
| uknp 2.4.1 | Retroviral | 2 | 16.94 | 34.02 | 2.01 |
| uknp 3.2.1 | Retroviral | 3 | 7.30 | 40.99 | 5.62 |
| uknp 3.2.2 | Retroviral | 3 | 32.75 | 191.18 | 5.84 |
| uknp 4.1.1 | Retroviral | 4 | 17.11 | 96.81 | 5.66 |
| uknp 5.2.1 | Retroviral | 5 | 13.38 | 128.37 | 9.59 |
| uknp 6.1.2 | Retroviral | 6 | 11.04 | 20.17 | 1.83 |
| 168_Cl T/F | Retroviral | 1b | 85.14 | 360.00 | 4.23 |
| 277_Cl T/F | Retroviral | 3a | 4.60 | 3.30 | 0.72 |
| 360_Cl T/F | Retroviral | 1a/2b | 3.90 | 35.30 | 9.05 |
| 686_Cl T/F | Retroviral | 1a | 6.90 | 25.50 | 3.70 |
| 4032_Cl T/F | Retroviral | 3a | 3.90 | 1.70 | 0.44 |
| 4087_Cl T/F | Retroviral | 1b | 0.00 | 128.00 | 128.00 |
| 023_Ch T/F1 | Retroviral | 1a | 12.13 | 64.50 | 5.32 |
| 023_Ch T/F2 | Retroviral | 1a | 0.00 | 10.30 | 10.30 |
| 023_197DPI | Retroviral | 1a | 0.00 | 5.30 | 5.30 |
| 240_Ch T/F | Retroviral | 3a | 3.00 | 2.30 | 0.77 |
| 256_Ch V1 | Retroviral | 1a | 9.70 | 88.00 | 9.07 |
| 256_Ch V2 | Retroviral | 3a | 6.00 | 97.00 | 16.17 |
| 256_79DPI 1 | Retroviral | 3a | 0.00 | 4.00 | 4.00 |
| 256_287DPI | Retroviral | 3a | 3.00 | 2.00 | 0.67 |
| HOK_Ch T/F | Retroviral | 1b | 12.90 | 155.00 | 12.02 |
| HOK_30DPI | Retroviral | 1b | 4.00 | 166.00 | 41.50 |
| HOK_233DPI | Retroviral | 1b | 0.00 | 5.70 | 5.70 |
| THD T/F | Retroviral | 1a | 14.30 | 71.30 | 4.99 |
| THD_109DPI | Retroviral | 1a | 114.30 | 74.50 | 0.65 |
| THD_198DPI | Retroviral | 1a | 0.00 | 104.00 | 104.00 |
| THG_Ch T/F | Retroviral | 1a | 14.30 | 71.30 | 4.99 |
| THG_58DPI | Retroviral | 1a | 82.50 | 64.80 | 0.79 |
| THG_184DPI | Retroviral | 1a | 47.00 | 49.00 | 1.04 |

**Supplementary Table 1. Signal-to-noise ratio (S/N) for all HCV strains tested in the study**. [a] E1E2 identification, [b] gag-pol genes origins: Lentiviral (HIV) and Retroviral (MLV), [c] fold change in HCVpp S/N (relative to S/N of parental 293T produced HCVpp), T/F: Transmission founder, ✤ E1E2 sequence of H77 mutants generated via site-directed mutagenesis, ◆ E1E2 sequence of an unpublished full-length cell-culture adapted J6/JFH chimeric strain.
